# First record of the subfamily Eucerotinae (Hymenoptera: Ichneumonidae) from the mainland Afrotropics, with a description of a new species

**DOI:** 10.64898/2026.05.11.724332

**Authors:** Tapani Hopkins, Alexsandra Nascimento, Bernardo F. Santos, Tomáš Hovorka, Ilari E. Sääksjärvi, Emil M. Österman

## Abstract

The ichneumonid subfamily Eucerotinae has been thought to be almost absent from the tropics, with the only known Afrotropical species found in Madagascar. We report the subfamily to be present in the mainland Afrotropics, and describe a new species, *Euceros species 1* from Uganda and Cameroon (name not yet shown in preprint). The subfamily had likely not been observed in the mainland Afrotropics before due to low abundances and insufficient sampling. More Eucerotinae likely remain to be discovered in tropical Africa and Asia, although tropical America may genuinely have few eucerotine species. Much more extensive sampling will be needed before it is possible to make confident estimates of how eucerotine diversity is distributed globally.

## Introduction

Eucerotinae is a small subfamily of Ichneumonidae (Hymenoptera) consisting of 51 species in the genus *Euceros* and one species in the genus *Barronia* (Gauld and Wahl 2002, Riedel 2018, 2025, Dal Pos et al. 2026). The subfamily’s known distribution is mainly from outside the tropics, especially from the northern temperate regions and temperate Australia (Gauld and Wahl 2002, Dal Pos et al. 2026). Only a single species of Eucerotinae has been recorded from tropical America, *E. limatus* Barron, 1978 from southeastern Brazil (Barron 1978, Dal Pos et al. 2026). South America also has a non-tropical species, *Barronia araucaria* Gauld & Wahl, 2002 from Chile (Gauld and Wahl 2002, Dal Pos et al. 2026), and two eucerotines collected in 1964 could be a further Chilean species if they are not *Barronia* (Porter 1997, their current location is unknown). Only a single species is currently known from tropical Africa, *E. madecassus* Seyrig, 1934 from Madagascar (Dal Pos et al. 2026). The only other known African species is *Euceros tunetanus* (Schmiedeknecht, 1900) from North Africa (Barron 1978, Dal Pos et al. 2026). Tropical Asia has two species recently recorded from Laos, *E. kiushuensis* Uchida, 1958 and *E. rufocinctus* (Ashmead, 1906) (Riedel 2025), and some species like *E. latitarsus* Barron, 1978 whose distribution may stretch into the northernmost parts of tropical Asia (Dal Pos et al. 2026). Otherwise tropical Asia is devoid of Eucerotinae observations, although recent collecting in Thailand (Quicke et al. 2023) appears to have caught one Eucerotinae of a likely undescribed species (identifier THAMK10015-23 in Anon 2023, GBIF.org 2025a).

The biology of the subfamily is highly unusual, and as far as is known unique for Ichneumonidae, resembling that of the hymenopteran family Trigonalidae (Carmean and Kimsey 1998). Females lay stalked eggs on plants, from which the larvae attach themselves to passing caterpillars or sawfly larvae (Tripp 1961, Gauld and Wahl 2002, Shaw 2014). If or when the host is parasitised by some other ichneumonoid wasp, the eucerotine larva enters the ichneumonoid larva and develops inside it as a hyperparasitoid.

The systematic position of the subfamily is unclear. It has at different times been proposed to be related to, or even part of, subfamilies Lycorininae, Tryphoninae, Ctenopelmatinae, Metopinae, Microleptinae or Brachycyrtinae, but with little convincing evidence for any of these except Brachycyrtinae, with which it is currently believed to be related (Gauld and Wahl 2002, Quicke 2015: 582, Santos 2017, Broad et al. 2018).

In this work, we report the subfamily to be present in the Afrotropics in mainland Africa, in at least Uganda and Cameroon, and describe a new species of *Euceros*. We also discuss the subfamily’s biogeographical distribution.

## Materials and Methods

The Ugandan material comes from the “Uganda 2014-2015” collecting event (Hopkins et al. 2019a, 2019b). In that sampling effort, 34 Malaise traps were kept for a year by one of the authors (Tapani Hopkins) in Ugandan forest and farmland, collecting 876 roughly two-week samples of flying insects, with a total sampling effort of 382.4 trap months. Some Eucerotinae were separated from the samples and pinned during the processing of other subfamilies, but the majority of Eucerotinae remain unprocessed. The pinned Eucerotinae are databased in the Finnish Biodiversity Information Facility (https://laji.fi/en), and their data in its current form is also stored in the Zenodo repository (Hopkins et al. 2026).

The Cameroonian material consists of two females, one of which was collected by one of the authors (Tomáš Hovorka) during a two-year (2024-2026) collecting event in southern Cameroon, with twelve Malaise traps in secondary forest near the village of Ebogo. All except the last batch of samples (August 2025-January 2026) were checked for Eucerotinae. One female specimen was found, collected in trap M2 in a forest clearing between two forest edges. Examination of museum material revealed another, older, specimen in the Natural History Museum, London.

To obtain DNA barcoding sequences for our two new species, we bioinformatically mined genomic data generated from the sequencing of ultraconserved elements (UCEs, Faircloth et al. 2012) as part of an ongoing study on the phylogeny of Ichneumonidae (Santos et al. in prep). Sequences for the Folmer fragment of the COI gene (Folmer et al. 1994) were pulled from the assembled reads using the script assembly_match_contigs_to_barcodes from the Phyluce v1.5 pipeline (Faircloth 2016).

We took focus stacked photographs of the Ugandan holotype specimen with a Canon EOS 7 D Mark II digital camera attached to an Olympus SZX 16 stereomicroscope. The programs QuickPHOTO MICRO 3.1 and Deep Focus 3.4 (by PROMICRA) handled the taking of individual photographs, and Helicon Focus 7.7.5 (by Helicon Soft Ltd.) combined them into a focus stacked image.

We measured the wing length from the photographs used for focus stacking, using the program QuickPHOTO MICRO 3.1. We got other measurements by viewing through the microscope, or measuring from the focus stacked images. Wing length was from the base to the furthest part of the wing. For the width versus height of the face, width was the minimum distance between the eyes, and height the vertical distance from the margin of the clypeus to the insertion of the antennae.

To verify our African species was new, we compared it to photographs of the holotype of the geographically nearest known species, *Euceros madecassus* in Madagascar (MNHN and Hervé 2014), photographs of a paratype and non-type specimen of *E. madecassus* (Noort 2025), to a non-type specimen of *E. madecassus* from the Natural History Museum, London (NHMUK 010887094), to the morphological codings of Eucerotinae in Gauld and Wahl (2002), and to CO1 barcodes we found via GBIF (GBIF.org 2025b) or in the Barcode of Life Database (https://boldsystems.org/). The two most relevant CO1 barcodes found this way, of *E. madecassus*, came from the Figshare repository (Miraldo et al. 2024, described in Miraldo et al. 2025) and GenBank (accession number JF963331.1, Sayers et al. 2025, described in Quicke et al. 2012) and are included in the dataset of this paper (Hopkins et al. 2026). We used objective clustering (Meier et al. 2006, https://github.com/asrivathsan/obj_cluster) to compare sequence divergence of the CO1 barcodes.

Morphological terminology follows that of Broad et al. (2018) and Goulet et al. (1993). The subfamily and generic diagnoses are largely based on Gauld and Wahl (2002). We wrote the diagnoses following the style outlined by Borkent (2021): short, clear diagnoses, which make it easy to confirm the wasp you are holding belongs to the taxon in question.

## Results

### DNA barcoding

Objective clustering revealed that the Ugandan and Cameroonian specimens assigned here to *Euceros species 1* differ by 3.67% in their COI sequences. This value exceeds the threshold commonly applied in insect DNA barcoding studies for preliminary species delimitation (e.g. Hebert et al. 2003, Cheng et al. 2023). However, the available material does not provide additional evidence supporting the recognition of two distinct species. Morphological differences between the populations are minor and inconsistent, and the Cameroonian lineage is currently represented by only two partially damaged specimens. Given the limited sampling and absence of clear diagnostic morphological characters, we conservatively interpret the Ugandan and Cameroonian specimens as representing a single species pending the availability of additional material. Future sampling across mainland African populations may help clarify whether the observed COI divergence reflects intraspecific geographic structure or species-level differentiation.

The two specimens of *E. madecassus* differed by only 0.75% in their COI sequences, while differing over 10% from our newly described species. No other COI sequences available on BOLD had less than 8.28% divergence to our new species.

### Taxonomy

#### Subfamily Eucerotinae Viereck, 1919

Type genus: *Euceros* Gravenhorst, 1829

#### Diagnosis

*Female*: Ovipositor tiny and weakly sclerotised (c.f. figure 1C). Areolet open with no trace of wing vein 3rs-m, spiracle of tergite 1 anterior to centre, anterior tergites wide. Antennae often centrally slightly flattened and widened. *Male*: Areolet open with no trace of wing vein 3rs-m, spiracle of tergite 1 anterior to centre, anterior tergites wide. Antennae centrally flattened and/or widened, often strikingly so.

**Figure 1:**
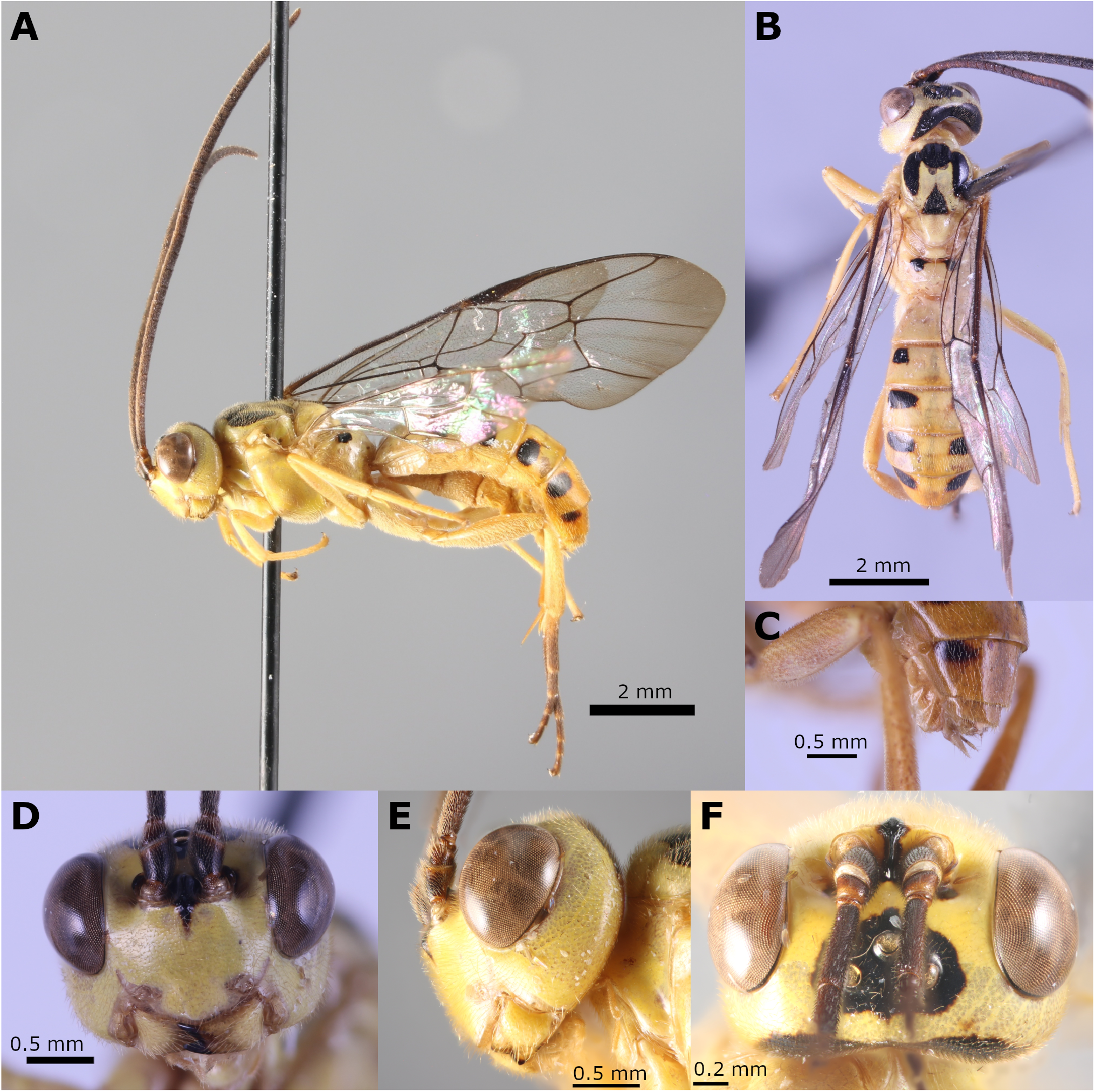
*Euceros species 1* holotype (http://mus.utu.fi/ZMUT.28283). **A**. habitus, lateral view **B**. habitus, dorsal view (picture from before the left fore wing came loose) **C**. tip of metasoma, including the ovipositor **D**. head, frontal view **E**. head, lateral view **F**. head, dorsal view.

#### Remarks

In practice, the unusual ovipositor of the females and the antennae of the males, together with the distinctive robust habitus (especially the stout tergites), make Eucerotinae relatively easy to recognise. The female ovipositor is so vestigial, they may be mistaken for males at a quick glance. We have found the pectinate hind claws of Eucerotinae to be a useful additional character: some African male Acaenitinae, for example, can be surprisingly easy to confuse with female Eucerotinae, but lack the pectinate claws.

#### Genus *Barronia* Gauld & Wahl, 2002

Type species: *Barronia araucaria* Gauld & Wahl, 2002

#### Diagnosis

Concave apical margin of clypeus (c.f. figure 6 Gauld and Wahl 2002), unlike *Euceros* where it is convex (c.f. 2B) or sometimes truncate. Clypeal sulcus between clypeus and face strongly impressed, unlike most *Euceros*.

#### Remarks

*Barronia* currently consists of just one species, *Barronia araucaria*, found in Chile. Gauld and Wahl (2002) appear to have chosen to give only the most relevant character states for telling the genus apart from *Euceros*, with their data matrix (Hopkins et al. 2026) having only two character states that *Barronia* does not share with at least one Euceros species (the convex apical margin of the clypeus, state 4 in the matrix, and the insertion of the hind coxa, state 38 in the matrix). However, the species looks distinctly different in photos of the holotype that we have seen.

#### Genus *Euceros* Gravenhorst, 1829

Type species: *Euceros crassicornis* Gravenhorst, 1829

#### Diagnosis

Convex or sometimes truncate apical margin of clypeus (c.f. 2B), unlike *Barronia* where it is concave (c.f. figure 6 Gauld and Wahl 2002). Clypeal sulcus between clypeus and face absent or weakly impressed in most species, unlike in *Barronia* where it is strongly impressed.

#### Remarks

Currently, all Eucerotinae except for the one Chilean species *Barronia araucaria* are in the genus *Euceros*.

#### *Euceros species 1* (name not yet shown in preprint)

Figures 1A-F, 2A-K, 3A-D

#### Material examined

Five females, basic data in Table 1 and complete data in the Zenodo repository (Hopkins et al. 2026).

**Table 1:**
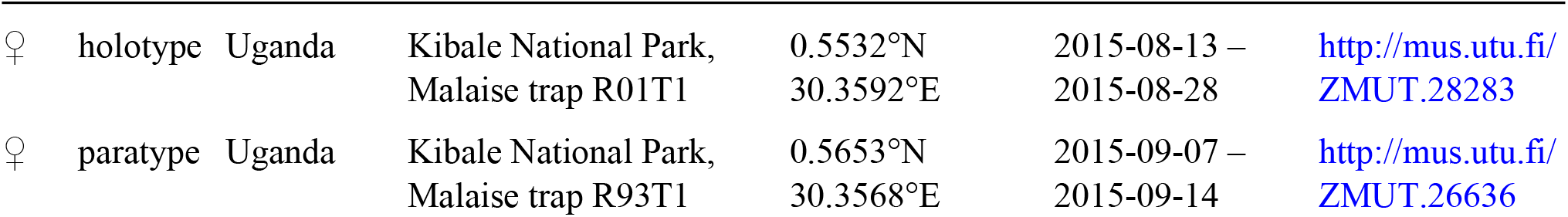
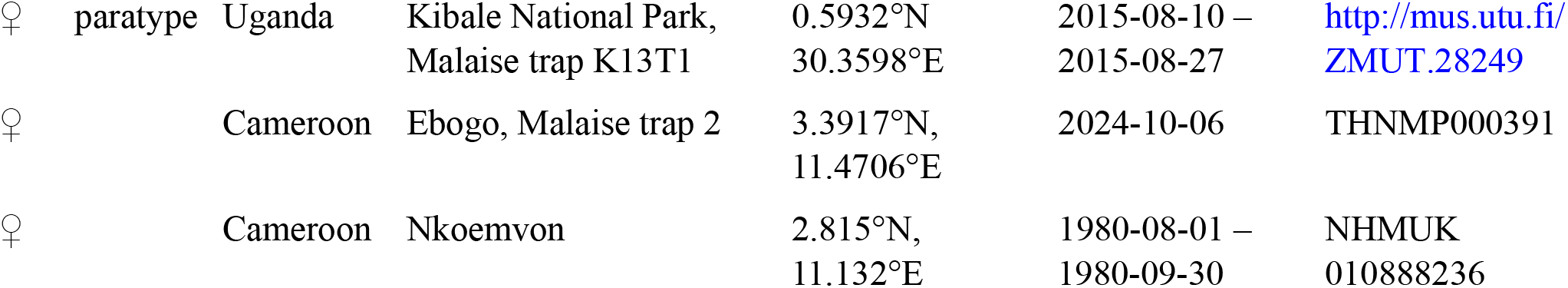
Material examined, *Euceros species 1*.

|  |  |  |  |  |  |  |
| --- | --- | --- | --- | --- | --- | --- |
| ♀ | holotype | Uganda | Kibale National Park,<br>Malaise trap R01T1 | 0.5532°N<br>30.3592°E | 2015-08-13 –<br>2015-08-28 | http://mus.utu.fi/<br>ZMUT.28283 |
| ♀ | paratype | Uganda | Kibale National Park,<br>Malaise trap R93T1 | 0.5653°N<br>30.3568°E | 2015-09-07 –<br>2015-09-14 | http://mus.utu.fi/<br>ZMUT.26636 |
| ♀ | paratype | Uganda | Kibale National Park,<br>Malaise trap K13T1 | 0.5932°N<br>30.3598°E | 2015-08-10 –<br>2015-08-27 | http://mus.utu.fi/<br>ZMUT.28249 |
| ♀ |  | Cameroon | Ebogo, Malaise trap 2 | 3.3917°N,<br>11.4706°E | 2024-10-06 | THNMP000391 |
| ♀ |  | Cameroon | Nkoemvon | 2.815°N,<br>11.132°E | 1980-08-01 –<br>1980-09-30 | NHMUK<br>010888236 |

#### Diagnosis

*Female*: Only known mainland Afrotropical *Euceros* species. The weak or absent posterior transverse carina of the propodeum and the lateral black spots of the metasoma may be useful additional characters (but see Remarks). *Male*: unknown, but likely to share the same diagnostic characters, since other *Euceros* species are not known to differ strongly between sexes other than in the antennae.

#### COI barcode

Paratype http://mus.utu.fi/ZMUT.28249:

acaaatcacaaagatattggtattttatattttattttcggtttatgatcaggtctattaggatcttctctaagagtattaattcgtttagaattaggta atcctgggtatttaattaataatgatcaaatttataactctatagtaacaattcatgcttttgtaataattttttttttagttatacctgtaataattgg gggatttggaaattgattagttcctttaataattggggctcctgatatagcttttcctcgtataaataatataagattttgacttttaattccttctata ttattattattatctagaagattttctaaccaaggaataggaactggatgaacggtttatcctcctttatctttaaatataagacatgagggtatatgtg tagatttgggaattttttctttacatttagcaggtatatcttctattataggagcaattaattttattactacaattttaaatataacac

Cameroonian specimen THNMP000391:

acaaatcataaagatattggtattttatattttattttcggtttatgatcaggtttattgggatcttctttaagagtattaattcgattagaattaggta atcctgggtatttaattaataatgatcaaatttataattctatagtaacaattcatgcttttgtaataattttttttttagttatgcctgtaataattgg aggatttggaaattggttagttcctttgataattggggctcctgatatagctttccctcgtataaataatataagattttgacttttaattccttctata ttattattattatctagaagattttctaatcaaggaataggaactgggtgaacggtttatcctcctttatctttaaatataagacatgaaggtatatgtg tagatttaggaattttttccttacatttagcaggtatatcttcaattataggggcaattaattttattactacaattttaaatataacaccatataatat aaaatatgaacaattacctttatttgtatgatctattaatattacagctattttattattattagcagttcctgtattagctggtgctattactatatta ttaacagatcgaaacttaaacacttctttttttgatccttcaggaggaggggatcctattttatatcaacatttattttgattttttggtcaccccgaa

#### Description

Holotype female http://mus.utu.fi/ZMUT.28283. Fore wing length 8.7 mm.

##### Head

Mandibles stout, with dorsal tooth slightly longer than ventral. Margin of clypeus convex when viewed from the front (figure 2B). Clypeal sulcus between clypeus and face not impressed, with no clear border between the clypeus and the rest of the face (figure 2B). Face about 1.3–1.4 times as wide as high (figure 2B). Occipital carina and hypostomal carina reach the base of the mandibles without merging (figure 2C). Ridge-like carina in between antennal sockets (figure 2D). Antennae with 33 flagellomeres. Antennae very slightly flattened and broadened near centre.

**Figure 2:**
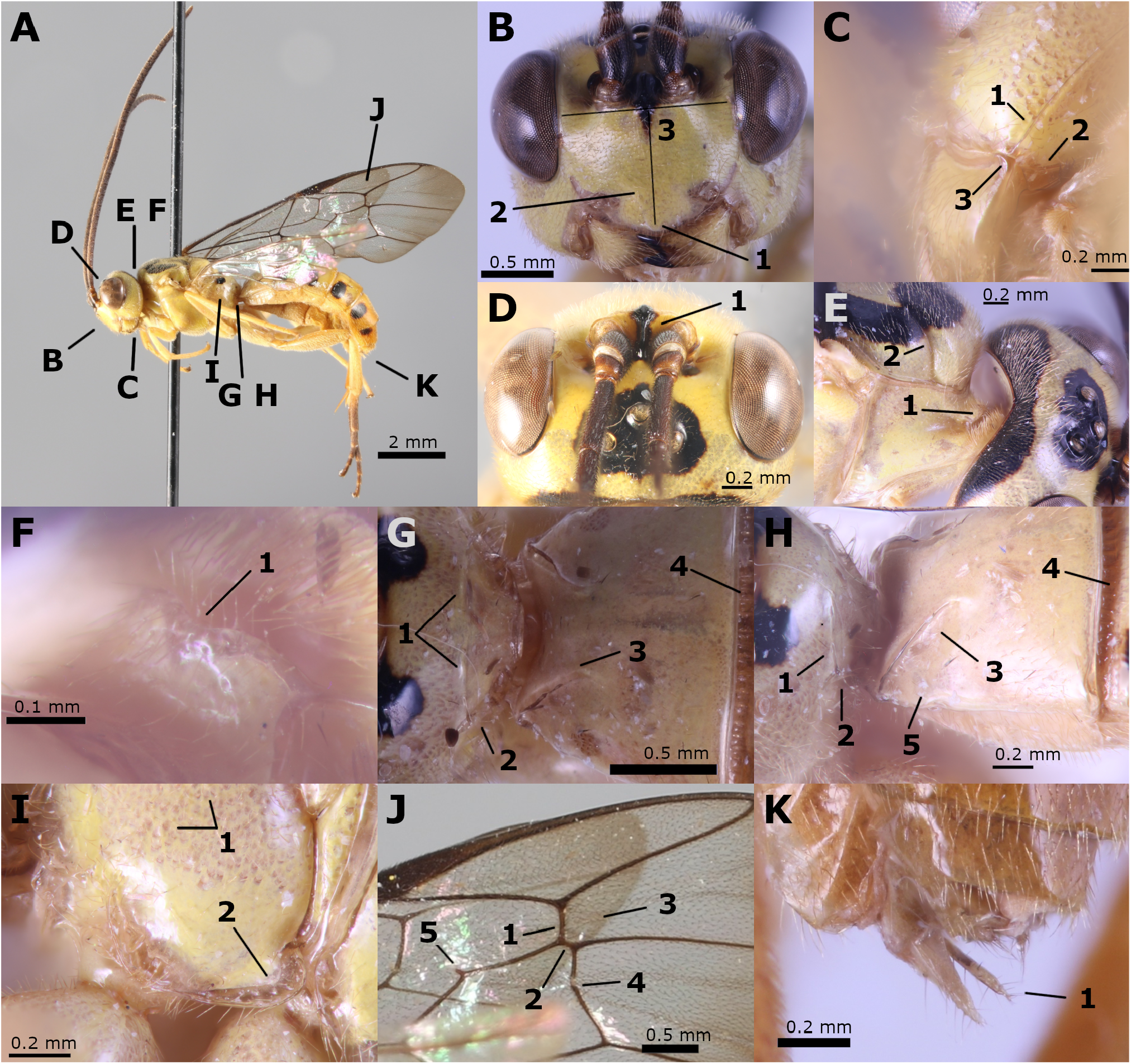
Characters of *Euceros species 1*. All images are of the holotype (http://mus.utu.fi/ZMUT.28283). **A**. habitus in lateral view, showing where panels B-K are from **B**. head in frontal view, showing the convex clypeus margin (1), lack of a clear clypeal sulcus (2), and the width and height of the face (3) **C**. lower head in lateral view, showing that the occipital carina (1) and the hypostomal carina (2) do not merge before the base of the mandible (3) **D**. head in dorsal view, showing the ridge between the antennal sockets (1) **E**. neck, showing the pronotal lobe (1) and the notauli (2) **F**. pronotum lobe, showing the mediodorsally concave edge (1) **G**. propodeum and tergite 1 in dorsal view, showing the weak posterior transverse carina (1), the short lateral longitudinal carina (2), the lateromedian carina of tergite 1 (3), and the anterior transverse groove of tergite 2 (4) **H**. propodeum and tergite 1 in lateral view, showing the weak posterior transverse carina (1), the short lateral longitudinal carina (2), the lateromedian carina of tergite 1 (3), the anterior transverse groove of tergite 2 (4), and the spiracle of tergite 1 (5) **I**. propodeum and metapleuron in lateral view, showing the lack of a pleural carina (1), and the submetapeural carina anteriorly expanded into a lobe (2) **J**. fore wing, showing that 2rs-m (1) is much longer than the segment (2) between it and 2m-cu, the lack of 3rs-m (3), the one bulla in 2m-cu (4), and the ramulus (5) **K**. tip of metasoma, showing the small weakly sclerotised ovipositor (1).

##### Mesosoma

Pronotum with a mediodorsal lobe (figure 2E), the apex of which has a concave notch (figure 2F). Epomia absent. Mesoscutum smooth with fine hairs, with strongly impressed notauli (figure 2E). Mesopleuron smooth, with sternaulus absent and with epicnemial carina reaching above level of lower corner of pronotum. Propodeum smooth; anterior transverse carina absent; posterior transverse carina present in between lateral longitudinal carinae, but very weak (figure 2G, H); lateral longitudinal carinae present posterior to the posterior transverse carina, absent otherwise (figure 2G, H). Pleural carina absent, so there is no clear border between the propodeum and metapleuron (figure 2I). Submetapleural carina anteriorly expanded into a rounded lobe (figure 2I). Legs with tarsal claws pectinate.

##### Wings

Fore wing with 2rs-m long and very close to 2m-cu, so that the segment of M between the two veins is much shorter than 2rs-m (figure 2J). 3rs-m absent (figure 2J). 2m-cu with one bulla (figure 2J). 1m-cu&M with a very short ramulus (figure 2J).

##### Metasoma

Spiracle of tergite 1 anterior to centre (figure 2H). Tergite 1 smooth, with lateromedian carinae stretching from the anterior corners diagonally towards the middle, to about 0.4 times the length of the tergite (figure 2G, H). Tergite 2 with an anterior transverse groove (figure 2G, H), similar grooves on at least tergites 3–4, although not as easily visible due to being covered by preceding tergite. Ovipositor very small and weakly sclerotised (figure 2K).

##### Colour

General colour yellow. Black mandibular teeth, area between antennal sockets, interocellar area, vertex, two median spots anterior and posterior on mesoscutum and two lateral stripes on mesoscutum, two lateral spots on anterior of propodeum, lateral spots on tergites 2–6. Antennae dark brown. Hind tarsi dark brown. Wings hyaline but with denser hairs making the forewings slightly darker apically.

#### Remarks

In practice, *E. species 1* is easy to recognise since there are only two other known African species, of which *E. madecassus* is likely confined to Madagascar and *E. tunetanus*, from outside the Afrotropics in North Africa, is mostly black (Barron 1978). In the material we have examined, *E. species 1* also differs from *E. madecassus* in the lateral black spots of the metasoma: the spots are present from tergite 2 onwards in the Ugandan material and absent in the Cameroonian material, whereas *E. madecassus* has the spots from tergite 3 onwards. The strength of the posterior transverse carina of the propodeum may also distinguish the two species. However, when comparing specimens to *E. madecassus* it should be noted that some of the available information about that species is contradictory. Gauld and Wahl (2002) claim it has vestigial notauli and that the submetapleural carina is not expanded into a lobe, and a photograph of the holotype shows no submetapleural lobe (MNHN and Hervé 2014); yet notauli are clearly visible in photographs of the holotype and other specimens (MNHN and Hervé 2014, Noort 2025), photographs of other specimens seem to show a submetapleural lobe (Noort 2025), and the non-type specimen we examined has a submetapleural lobe. These may just be mistakes in the data or unclear photographs, but we have a sneaking suspicion there may be several (likely undescribed) species involved, which are getting misidentified as *E. madecassus*.

The Ugandan and Cameroonian specimens of *E. species 1* have quite a large difference in their COI barcode (3.67%), and milder differences in morphology and colour. We suspect they may actually be two separate species, but do not feel confident enough to split the species until more material can be examined.

#### Variation

The posterior transverse carina of the propodeum is missing in some specimens. The number of flagellomeres varies from 31 to 33. The fore wing length of the Cameroonian specimen THNMP000391 is 9.4 mm (c.f. holotype 8.7 mm, other specimens not measured). The ramulus in the fore wing varies from vestigial to absent.

The Ugandan specimens have a clearer concave notch on the mediodorsal lobe of the pronotum than the Cameroonian specimens, whose lobe is very shallowly incised, if at all. However, this character is very hard to see accurately on some of the specimens.

The general colour varies from bright yellow (the holotype), to faded whitish yellow with the tip of the metasoma bright yellow (paratypes), to dark yellow (Cameroonian specimens, c.f. figure 3). The head of one Cameroonian specimen (THNMP000391) is a burnt ochre in colour, the other bright yellow. We suspect much of this colour variation may be due to fading or preservation (e.g. THNMP000391 had the inner tissues digested for DNA extraction), and that the the holotype is closest to the natural colour.

**Figure 3:**
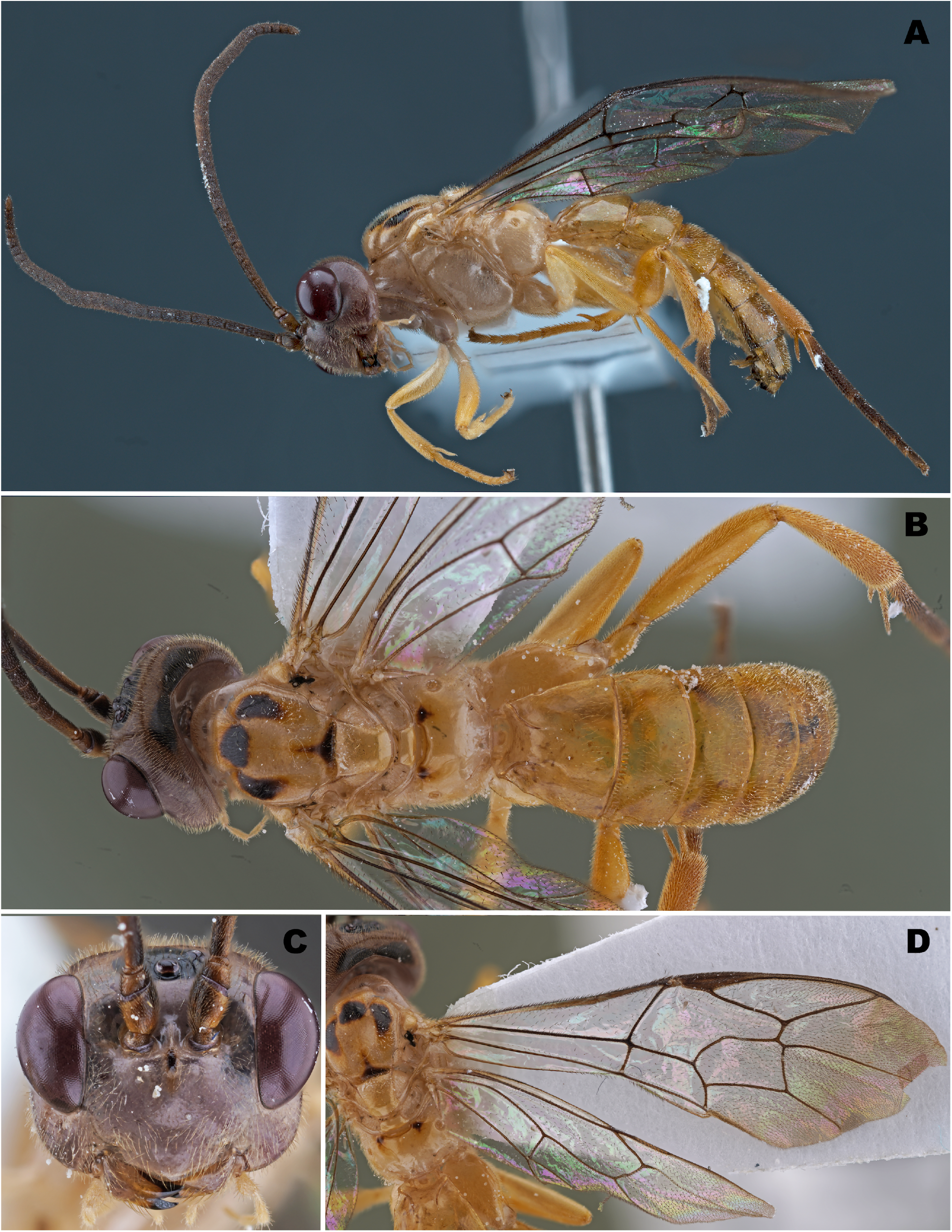
*Euceros species 1* Cameroonian specimen THNMP000391. **A**. habitus, lateral view **B**. habitus, dorsal view **C**. head, frontal view **D**. wings.

The Ugandan specimens have lateral black spots on tergites 2 onwards (figure 1B), while the Cameroonian specimens lack these spots (figure 3B).

#### Etymology

This species has not yet got a name for the preprint. We intend to ask readers of the local newspaper, Österbottens Tidning, to send in name suggestions and vote for a good name.

#### Holotype condition

The left fore wing of the holotype is loose.

## Discussion

It is obvious that the subfamily Eucerotinae, despite earlier assumptions to the contrary (Gauld and Wahl 2002), is widespread in mainland tropical Africa. We found a new species in tropical forest in both Uganda and Cameroon; it is exceedingly unlikely that the subfamily wouldn’t be present in, for example, the Democratic Republic of the Congo, in between these two countries. The mainland Afrotropical species (which may in fact be two separate species) is clearly closely related to the Malagasy species, and the ancestors of the Afrotropical species have presumably either spread from Madagascar to the mainland or vice versa. If Gauld and Wahl (2002) are correct that eucerotines have a Gondwanan origin and spread to the northern hemisphere from the Malagasy/Indian tectonic plate, a Malagasy origin is more likely. However, assumptions of a Gondwanan origin for both Eucerotinae and the possibly related subfamily Brachycyrtinae were partly based on the belief they are absent from the mainland Afrotropics (Gauld and Wahl 2002), which we now know to be incorrect for both subfamilies (cf. Giovanni and Varga 2021 for Brachycyrtinae).

The lack of earlier observations of this subfamily from the mainland Afrotropics is likely due to its low abundances and insufficient sampling. It will be many years before we know how common the new species is in our Ugandan material, but a rough estimate based on what little material has been processed suggests tens of individuals rather than hundreds in total, collected in 876 samples over a whole year. As a comparison, some of our common Ugandan species (e.g. *Echthromorpha agrestoria* (Swederus, 1787)) number over a thousand individuals. Collecting in non-tropical Georgia, USA also caught very low abundances of Eucerotinae (Gaasch et al. 1998), nor do we know of any material where Eucerotinae would be abundant. Low abundances are likely the norm for this subfamily throughout its distributional range, perhaps as a result of the family’s highly specialised biology. Such low abundances simply cannot be expected to be detected with the smaller-scale sampling which has largely been the norm in Africa up to now (e.g. Noort et al. 2000, Noort 2004 which caught no Eucerotinae).

At a global scale, the presumed near-absence of the subfamily Eucerotinae in the tropics (Gauld and Wahl 2002) seems doubtful. The known diversity of the subfamily is still strongly focused on the northern temperate regions and temperate Australia, but we have discovered a species in tropical mainland Africa, and an undescribed species appears to have recently been found in tropical Asia (Quicke et al. 2023, identifier THAMK10015-23 in Anon 2023). More extensive collecting with numerous Malaise traps played a major part in both these discoveries. It thus seems likely that even more species remain undiscovered in both tropical Africa and Asia, and that further collecting is needed before we can tell what the true eucerotine diversity of these regions is. The case for a near-absence of eucerotines in tropical America, on the other hand, is stronger. The only known tropical American species is from southeastern Brazil in high altitude Atlantic forest (20°S: Barron 1978, Dal Pos et al. 2026). No eucerotines were caught in the Peruvian and Ecuadorian Amazon, in extensive collecting with Malaise traps and canopy fogging that matched our Ugandan collecting in intensity (Sääksjärvi et al. 2004, Gómez et al. 2017, summarised in Veijalainen et al. 2013, Hopkins et al. 2024), the material of which is at the Zoological Museum of the University of Turku. Nor did Gauld (1991) report the subfamily Eucerotinae in Costa Rica, despite one of the most intense tropical Malaise trapping programmes ever conducted. However, we do not feel the current evidence rules out the possibility of further collecting discovering more tropical American species, merely that a diverse eucerotine fauna seems implausible. The overall picture suggested by the new tropical discoveries is that while it is plausible that the subfamily Eucerotinae is more diverse in the temperate areas than the tropics, there is currently too little data to make confident estimates of its global distribution.

Although there is not enough data, the currently most plausible biogeographic distribution for Eucerotinae is for there to be the most species in northern temperate regions and temperate Australia, fewer species in tropical Asia and Africa, and almost no species in tropical America. If further collecting confirms such a pattern, we speculate that part of the reason for it, beyond the obvious host availability (e.g., few sawfly species in many tropical regions), could be one of predation pressure or competition from the biologically similar Trigonalidae. The unusual biology of eucerotines, with larvae exposed on leaves while waiting for their host, could make them sensitive to predation by e.g. ants, which would drop abundances and hence species richness in high predation environments. Predation pressure is generally highest in the tropics at low elevations (Roslin et al. 2017), which is exactly the kind of habitat in western Amazonia in which we have failed to discover eucerotines. Another, alternative, speculation is that the subfamilies Eucerotinae and Trigonalidae (which both have similar biologies) compete for the same hosts, and that Trigonalidae outcompetes Eucerotinae in the tropics. This would match the currently known distribution of Trigonalidae diversity, which is mainly tropical and subtropical (Carmean and Kimsey 1998).

## Acknowledgements

The collecting in Uganda was supported by Heikki Roininen, Isaiah Mwesige and the staff of the Makerere University Biological Field Station. Countless people helped process the Ugandan samples, including the staff of the Zoological Museum of the University of Turku, students of the university and school pupils from the Turku region. These contributions are gratefully acknowledged.

The required Ugandan research and export permits were issued by the Uganda National Council of Science and Technology (NS 504) and the Uganda Wildlife Authority.

## Data availability

The data of the wasps we examined is in the Zenodo repository (Hopkins et al. 2026, https://doi.org/10.5281/zenodo.15494332). The repository also has the CO1 barcode sequences we used, and the character state data matrix of Gauld and Wahl (2002) updated with the data of our new *Euceros* species.

